# Cortical spheroids show strain-dependent cell viability loss and neurite disruption following sustained compression injury

**DOI:** 10.1101/2023.11.15.567286

**Authors:** Rafael D. González-Cruz, Yang Wan, Dominick Calvao, Amina Burgess, William Renken, Francesca Vecchio, Christian Franck, Haneesh Kesari, Diane Hoffman-Kim

## Abstract

Sustained compressive injury (SCI) in the brain is observed in numerous injury and pathological scenarios, including tumors, ischemic stroke, and traumatic brain injury-related tissue swelling. Sustained compressive injury is characterized by tissue loading over time, and currently, there are few *in vitro* models suitable to study neural cell responses to strain-dependent sustained compressive injury. Here, we present an *in vitro* model of sustained compressive neural injury via centrifugation. Spheroids were made from neonatal rat cortical cells seeded at 4000 cells/spheroid and cultured for 14 days *in vitro*. A subset of spheroids was centrifuged at 209 or 419 rad/s for 2 minutes. Modeling the physical compression of the spheroids via finite element analyses, we found that spheroids centrifuged at 209 and 419 rad/s experienced pressures of 38 kPa and 149 kPa, respectively, and compressive strains of 18% and 35%, respectively. Quantification of LIVE-DEAD assay and Hoechst 33342 nuclear staining showed that centrifuged spheroids exhibited significantly higher DNA damage than control spheroids at 2, 8, and 24 hours post-injury. Immunohistochemistry of β_3_-tubulin networks at 2, 8, and 24 hours post-centrifugation injury showed increasing degradation of microtubules over time with increasing compressive strain. Our findings show that cellular injuries occur as a result of specific levels and timings of sustained compressive tissue strains. This experimental compressive injury model provides an *in vitro* platform to examine cellular injury to gain insights into brain injury that could be targeted with therapeutic strategies.

## Introduction

Sustained compression injury in the brain is a feature of many brain diseases and injuries (1,2). Increased intracranial pressure (ICP) exerted from expanding brain tumors, increased hematoma edema volume, and/or swollen injured tissue (3) can injure neighboring tissues and their cells by deforming them. While this pressure deforms adjacent brain tissues at generally lower strain magnitudes and rates than those observed in blunt and blast head trauma, the generated sustained compression injury (SCI) pressures typically last longer, ranging anywhere from minutes to months (4). Prolonged pressures can result in damage at the cellular level and, eventually, lead to altered electrophysiology, chronic neuroinflammation, and neuronal death. However, there is neither a clear understanding of the magnitude of the resulting deformations caused by these pressures nor of their durations. This knowledge gap represents a hindrance to understanding SCI progression and developing clinical strategies to prevent long-term brain damage caused by sustained compression.

Most compressive brain injury studies, either *in vivo* or *in vitro*, focus on replicating traumatic brain injury conditions by applying forces for very short durations at high speeds to mimic varying real-life injury scenarios. Such studies have allowed scientists to gain insight into the events following injuries, from blunt impact to blast-induced injury, as a function of strain and strain rate. For example, *in vivo* studies employing setups such as controlled cortical impact injuries on rats have allowed for monitoring the progression of neuronal disruption (5) and neuroinflammation (6,7) as well as injury biomarker release in live animals subjected to TBI (8). Meanwhile, *in vitro* experiments have allowed monitoring of live cell responses to compressive injury to determine injury thresholds based on strain magnitude and strain rate in both two- and three-dimensional culture platforms (9–13). While these studies provide information into the progression of strain-dependent brain injury, they typically employ quick loading and unloading conditions, and do not represent SCI.

Previous studies specific to SCI have been conducted *in vivo* using animal models and *in vitro* using culture platforms. *In vivo* SCI models include using setups similar to controlled cortical impact but delaying unloading times to minutes, rather than milliseconds (14). and implantation of a hemispherical plastic bead inside rat somatosensory cortices to compress the brain statically (15). *In vitro* SCI studies have typically applied pressurized air to 2D cultures of neural cell lines in pressurized culture chambers (16,17). These studies have begun to provide a time window into cellular events that occur after the exposure of the brain and neuronal cells to forces. However, there is a need for *in vitro* models that can achieve the combination of (1) controlling the pressures required to introduce compressive strains; (2) monitoring the resulting effects over time; and (3) doing so in a robust 3D model that recapitulates key features of the brain.

In this study, we present a new in *vitro injury* model as a platform to study sustained compression injury. We used a three-dimensional in vitro cortical spheroid (18–20) which mimics multiple characteristics of the *in vivo* brain that are highly relevant to injury response. Generated from postnatal rat cortical cells by self-assembly, the cortical spheroid contains interconnected cell types, cell density, and tissue stiffness of the *in vivo* cortex. Spheroid neurons are electrically active, with mature synapses and neurite networks in the first two weeks of culture. Spheroid cells include neurons, astrocytes, oligodendrocytes, microglia, and endothelial cells, and produce extracellular matrix, myelin, as well as capillary-like networks. All these characteristics are reproducible, and one postnatal rat cortex yields hundreds of spheroids, making this model an excellent choice for high throughput testing.

Here, cortical spheroids are deformed by pressure generated via centrifugation. The pressures are modulated by varying the centrifugation speeds and computed via finite element analysis (FEA). We demonstrate that sustained compression injury caused cellular death and neuronal network disruption in cortical spheroids within the first 24 hours following compression injury. Mechanics modeling and FEA allowed for the estimation of strains associated with the centrifugation-induced pressures and the concomitant changes observed in cell viability and neurite network organization. These results suggest that this model can facilitate the study of the effects of sustained compression and its resulting strains on the brain at the tissue and cellular levels.

## Materials and Methods

### Cortical dissection and spheroid culture

Cortical spheroids were assembled and cultured as described elsewhere, with minor modifications^1^. Briefly, cortices were dissected from postnatal day P2 CD neonatal male and female rat pups (Charles River) following University approved IACUC protocols and placed in cold Hibernate A with B27 Plus Supplement (Invitrogen). Then they were minced into small pieces and transferred to a 2 mg/ml papain solution, which was composed of lyophilized papain powder and Hibernate A without calcium chloride (BrainBits, LLC) and pre-warmed to 30°C. Cortical digestion in papain lasted for 30 minutes, with 5-minute agitation intervals. Single cortical cell suspensions were obtained by filtering and centrifuging the digested cell suspension at 150 RPM and seeded at 4000 cells/well in 96-well agarose hydrogels (see Fig. 1(b)) to form cortical spheroids (see Fig. 1(a)) via self-assembly. The agarose hydrogels were made by pouring a volume of 2% w/v molten agarose onto silicone molds (#24-96-Small, MicroTissues Inc), transferred to a 24-well polystyrene plate (see Fig. 1(c)) (CELLTREAT), and equilibrated with complete cortical media for 48 hours at 37°C prior to cell seeding. The 2% w/v agarose solution was made by dissolving 2g of ultrapure agarose powder (Invitrogen) in 100 mL of 1X phosphate buffer saline, pH = 7.4 (Gibco) and heating the 2% agarose solution in a microwave for 1 minute to solubilize the agarose powder. Self-assembled spheroids were cultured for 14 days *in vitro* (DIV14) at 37°C in complete cortical media. Complete cortical media consisted of Neurobasal A+ (Invitrogen), 1x B27 Plus Supplement, 0.5 mM GlutaMAX, and 1x Pen/Strep (Invitrogen).

**Fig. 1:**
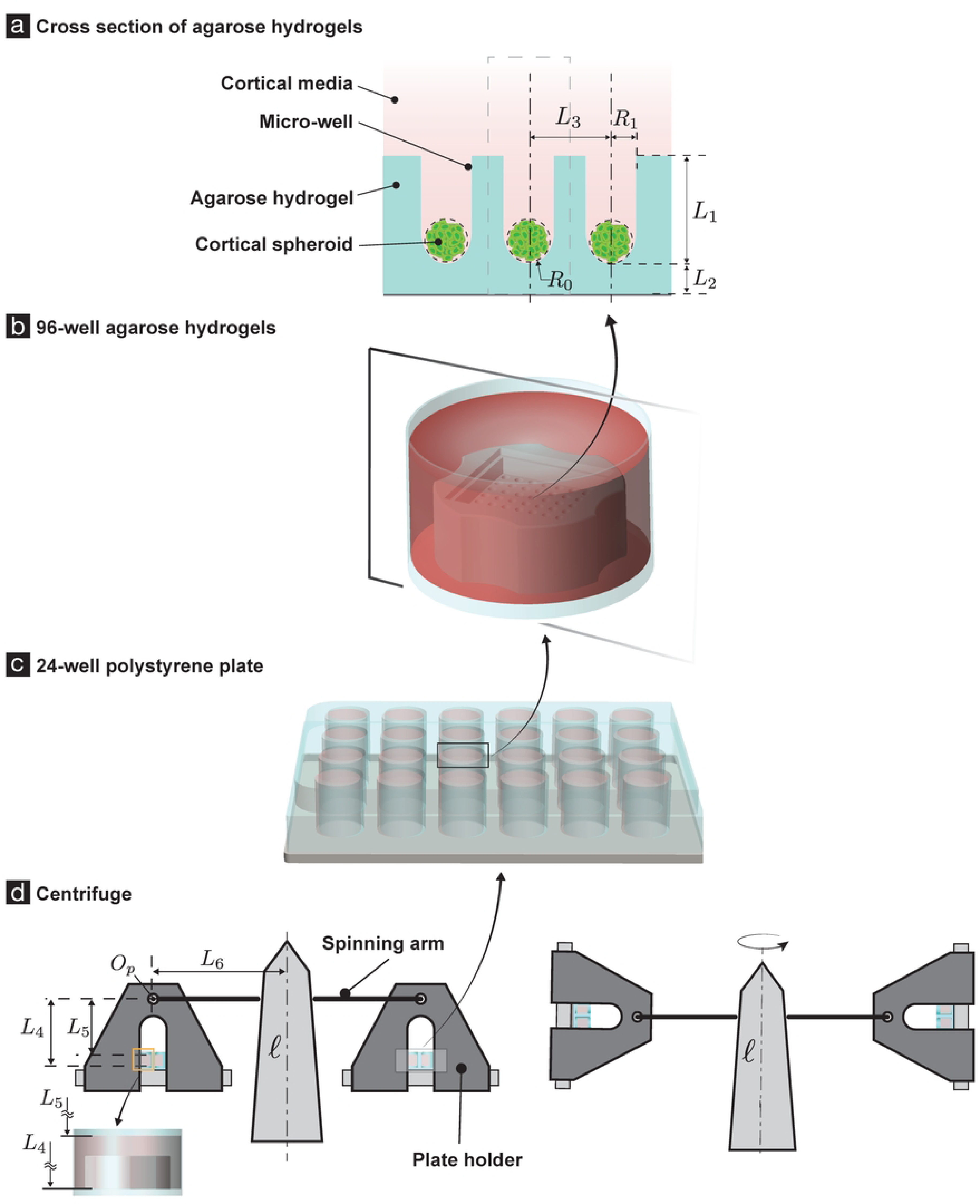
Experimental SCI setup and geometry of the spheroid centrifugation. DIV14 cortical spheroids were subjected to sustained compression injury via centrifugation. (a)–(d) shows the geometry of the spheroids’ centrifugation. The spheroids’ radii (𝑅_0_) varied from 75 to 85 microns. In our mechanics model (Fig. 2) we take the spheroid’s radii to be 80 microns. Each spheroid lies in a microwell that is 800 microns deep (𝐿_1_) and has a circular cross-section of radius 200 microns (𝑅_1_), with a hemispherical base. The micro-well is part of an agarose hydrogel structure (shown in cyan in (a)). The thickness of the agarose material under a spheroid is 1.5 mm (𝐿_2_). The agarose hydrogel contains 96 micro-wells in total, arranged in a square grid (b). The distance between adjacent micro-wells is 𝐿_3_, where 𝐿_3_ = 800 microns (a). The agarose hydrogel sits in one of the wells of a 24-well polystyrene plate (c). The polystyrene plate sits in a centrifuge cell culture plate holder (d), which is suspended from a centrifuge’s spinning arm. We refer to the point at which the centrifuge’s spinning arm attaches to the centrifuge cell culture plate holder as 𝑂_𝑃_ (d). We measured the base of the agarose hydrogel to be at a distance of 66 mm from the point 𝑂_𝑃_.(𝐿_4_). The spheroid and the microwell were submerged in cortical fluid media. From the dimensions of the polystyrene well, the volume of the agarose hydrogel structure, and the amount of fluid media added to a polystyrene well, we estimate that the surface of the cortical fluid media is at a distance of 59.61 mm from 𝑂_𝑃_. (𝐿_5_). The length of the spinning arm is 112.69 mm (𝐿_6_).

### Centrifugation

DIV14 cortical spheroids were subjected to sustained compression injury via centrifugation (Fig. 1). The spheroids were suspended in media in micro-wells of an agarose hydrogel structure (Fig. 1 (a)–(b)). The agarose hydrogel sat in one of the wells of a 24-well polystyrene plate (Fig. 1 (c)). Media was added to the wells of the polystyrene plate, so that the spheroids and the agarose-hydrogels were completely submerged by it. The size of the spheroid, the depth and thickness of the micro-well, the height of the fluid media in each polystyrene well as well as other important dimensions in the SCI experiment are described in Fig. 1’s caption. The polystyrene plate sat in a centrifuge cell culture plate holder (Fig. 1 (d)), which was suspended from a centrifuge’s spinning arm. The centrifuge was of tissue culture-grade (5810R Eppendorf). The spheroids were mechanically loaded (and as we shall show later, injured) by being spun by the centrifuge.

In each mechanical loading the angular velocity 𝜔 radians/seconds (rads/s) of the centrifuge was varied with time as:

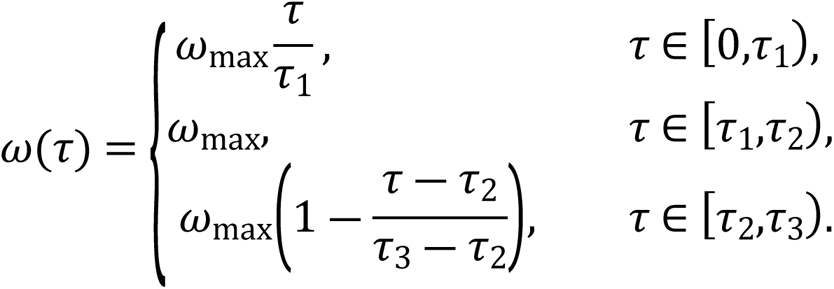

That is, the loading consisted of three stages: (St.1) an acceleration stage (from 0 seconds to 𝜏_1_ seconds) in which the angular velocity linearly increased from zero to 𝜔_max_ rads/s; (St.2) a constant angular velocity stage (from 𝜏_1_ seconds to 𝜏_2_ seconds), in which the angular velocity was kept constant at 𝜔_max_ rads/s, and; finally, (St.3) a deceleration stage (from 𝜏_2_ seconds to 𝜏_3_ seconds), in which the angular velocity decreased linearly from 𝜔_max_ rads/s to zero.

The spheroids were subjected to two different mechanical loadings; one in which 𝜔_max_ = 209 (𝜏_1_ = 10, 𝜏_2_ = 120, and 𝜏_3_ = 134), and another in which 𝜔_max_ = 419 (𝜏_1_ = 25, 𝜏_2_ = 120, and 𝜏_3_ = 144). These two angular velocities were chosen so that the whole range of speeds that our centrifuge was capable of accurately applying were explored uniformly. (Our centrifuge was advertised as being capable of a maximum angular velocity of 5000 RPM. However, we expected it to be capable of accurately applying speeds only up to 4000 RPM. Therefore, we choose the speeds of 0 RPM (control), 2000 RPM (209 rads/s), and 4000 (419 rads/s) RPM in our experiments.

### Mechanics model of the cortical spheroid, agarose hydrogel, and the cortical fluid media system

We constructed and solved a model that captures the mechanical interaction between the spheroid, the agarose hydrogel, and the cortical fluid media as they are spun by the centrifuge. The primary goal of the model was to estimate the strains and the pressures in the spheroid during the second stage of the loading (St.2). In our model we assume that in a frame that rotates with the centrifuge’s rotating arm all spatial mechanical fields are stationary with respect to time. The model was developed using the mechanics formalism of (21–23). The complete details of our model, including the various simplifying assumptions in it, are detailed in Supplementary Information (S1 Appendix) and (24).

Considering the spatial proximity of the cortical spheroids (see Fig. 1 (b)), while they are being centrifuged, we assume that all the spheroids experience the same order of magnitude of strains during the centrifugation. Consequently, we only model the deformation of a single cortical spheroid, along with the microwell containing it and the cortical media surrounding them. The geometry of the spheroid, the agarose hydrogel microwell, and the cortical fluid media in the reference configuration in our model is shown in Fig. 2 (a). The important dimensions in that geometry are described in Fig. 1, and their values are given in those figures’ captions.

**Fig. 2:**
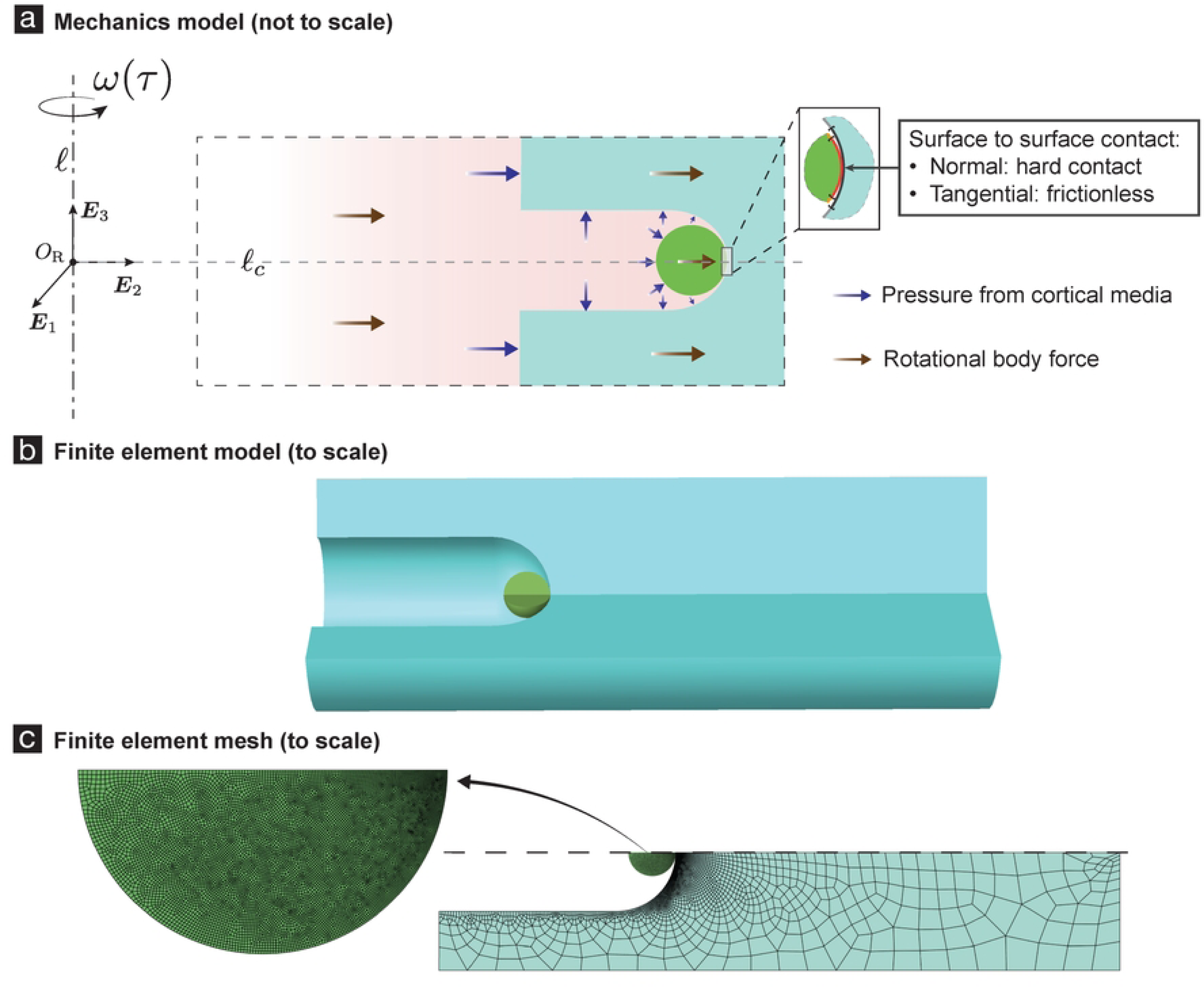
Mechanics model and finite element analysis of spheroid centrifugation (SCI experiments). (a) shows the geometry of our mechanics model of the SCI experiment. In our model we focus on a single spheroid. This is motivated by our assumption (a.1) that the stresses and strains in most of the spheroids are similar to each other (i.e., are of the same order of magnitude). We also assume (a.2) that the deformations and stresses in the spheroid and in the agarose hydrogel region in its vicinity are axi-symmetric. The spinning deforms and stresses the spheroid, and the agarose hydrogel, and creates additional pressures in the fluid media. We model this effect by introducing effective body forces (brown arrows in (a)) in each of them. We compute the pressure in the fluid media and use that to impose surface traction boundary conditions on the spheroid and the agarose hydrogel (purple arrows in (a)). We model the interaction between the spheroid and the agarose hydrogel using a non-adhesive, frictionless contact theory. The surface regions of the spheroid and the agarose hydrogel that eventually come into contact with each other are marked in red and dark-gray, respectively, in the inset of (a). **(b)** We use finite element (FE) techniques to solve the model in (a). The FE model contains a single spheroid and all the agarose hydrogel region that is at a distance of 𝐿_3_/2 microns from the central axis (𝓁_𝑐_) of the spheroid’s micro-well. We refer to this agarose hydrogel region as hydrogel cell (cyan in (b)). Recall that 𝐿_3_is the distance between two adjacent agarose hydrogel microwells (see Fig. 1 (a)). This feature of our FE model was motivated by our assumption (a.1). In the FE model we take all the mechanical fields to be axi-symmetric around the micro-well’s central axis. This feature is a consequence of our assumption (a.2). Also, as a consequence of this assumption the circumferential displacements in our FE model are naught. As a consequence of (a.1) and (a.2) we have that there are no radial displacements on the outer cylindrical surface of the hydrogel cell. We impose zero displacements at the base of the hydrogel cell. Other boundary conditions in our FE model are discussed in section *Centrifugation.* **(c)** shows a representative computational mesh of our FE model. The spheroid’s FE mesh typically contained round 11000 linear quadrilateral elements, and the hydrogel cell’s mesh typically contained around 3200 linear quadrilateral elements. After discretization and assembly, we solve around 29000 non-linear equations, using iterative numerical techniques.

We assume that the mechanics of the spheroid, the microwell composed of the agarose hydrogel, and the cortical fluid media can be well modeled using continuum mechanics theories. Consequently, we model the cortical spheroid and the agarose hydrogel microwell as homogenous solids, of densities 1240 mg/ml and 1640 mg/ml, respectively, and the cortical fluid media as a homogeneous fluid, of density 980 mg/ml. More specifically, we model the spheroid as a spherical ball composed of an incompressible neo-Hookean material of shear modulus 4/3 kPa; the micro-well as a structure composed of a compressible neo-Hookean material having shear modulus 107 kPa and bulk modulus 501 kPa; and the cortical media as an incompressible Newtonian fluid. It is unlikely that the spheroids have a purely elastic mechanical behavior, let alone the elastic behavior dictated by the incompressible neo-Hookean material model, as we assume here. However, since we do not have any direct and complete characterization of the spheroid’s mechanical behavior, we chose the incompressible neo-Hookean material model for the spheroid to simplify our analysis. We chose the value of 4/3 kPa for the spheroid’s shear modulus so that our model is consistent with previous studies data (25). Our choices for the agarose-hydrogel-microwell and the cortical-fluid-media’s material models are also based on similar reasoning.

The spheroid, the micro-well, and the fluid media all experience effective body forces due to the spinning, see Fig. 2 (a). We model their interactions with each other by applying to each of them suitable surface tractions and constraints on their deformations. In our model, we were able to explicitly compute the pressure field in the fluid media (see Section 4.3 in S1 Appendix and (24)). The interaction of the spheroid with the fluid media was captured by using that pressure field and constructing appropriate traction boundary conditions on the spheroid. The interaction of the microwell with the fluid media was captured similarly. The interaction between the spheroid and the microwell was modeled using non-adhesive frictionless contact boundary conditions. The other boundary conditions in our model are discussed in the caption of Fig. 2.

We set up boundary value problems (BVPs) for the spheroid and the microwell based on the above discussed body forces, surface tractions, and contact boundary conditions. Their BVPs turn out to be coupled due to the contact boundary conditions. We solve for the strains and the pressures in the spheroid by solving the coupled boundary value problems simultaneously using nonlinear finite element techniques (see Fig. 2 (b)). See S1 Appendix and (24) for a more detailed description of our mechanics model. Representative pressures and strains in the spheroid, as predicted by our model, are show in Fig. 3.

**Fig. 3:**
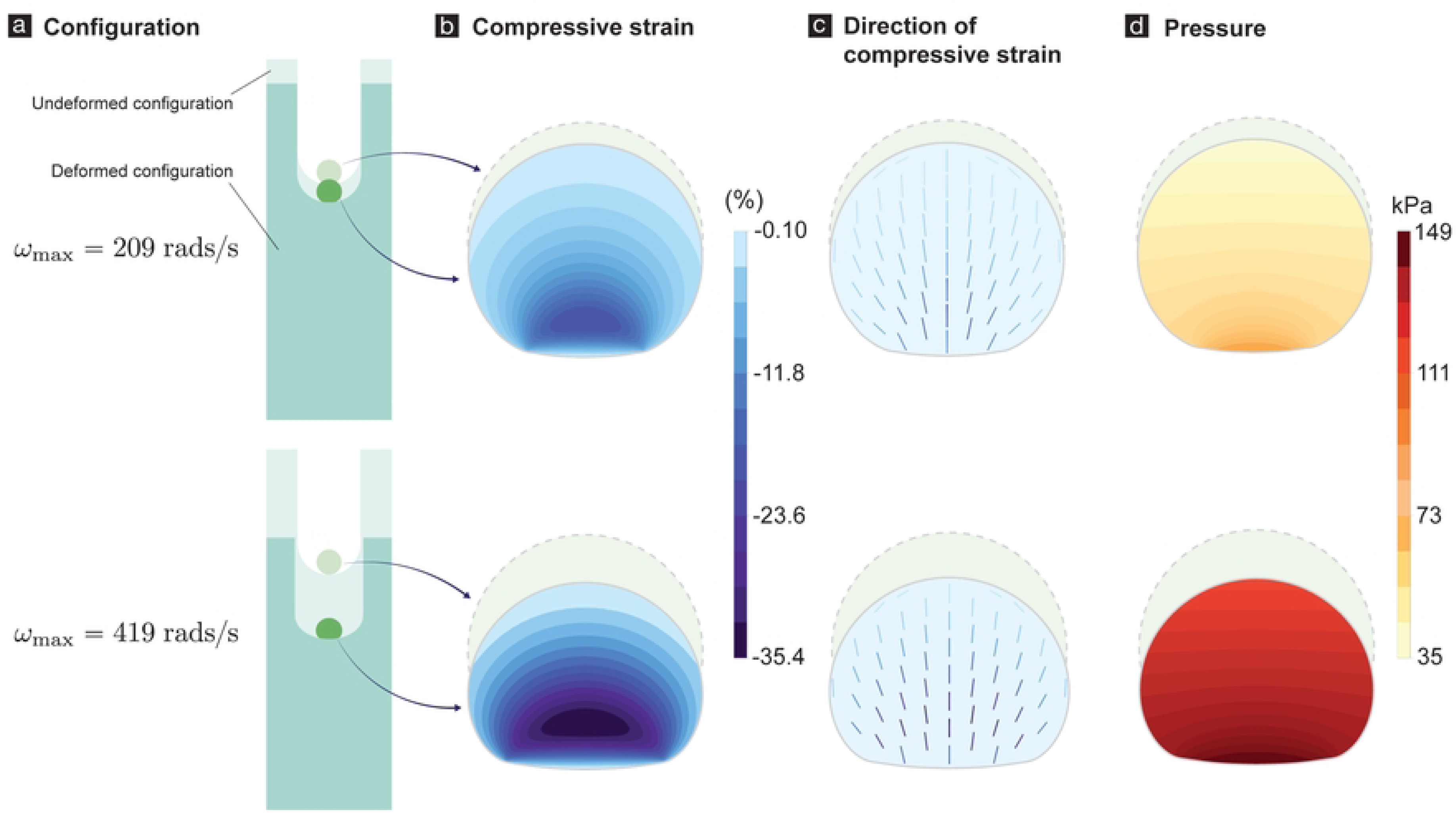
Deformations, strains, and pressures in a representative spheroid due to centrifugation at angular velocities of 209 rads/s (top row) and 419 rads/s (bottom row). (A) shows the undeformed and deformed configurations of the cortical spheroid and agarose hydrogel microwell. (B) shows the compressive strain, i.e., the minimum eigenvalue of the logarithmic strain tensor. For details on the logarithmic strain tensor see (24). (C) Each line segment shows a section of the fiber associated with the eigenvectors that correspond to the minimum eigen value of the logarithmic strain tensor. (D) shows the pressures in the spheroid.

### Sustained compression injury (SCI) experiments

To quantify cell death in centrifuged spheroids we incubated a subset of them for 0, 2, 8, and 24 hours after centrifugation and then stained them with 2µM ethidium homodimer 1 (EthD1) in complete cortical media and 20 µg/mL Hoechst (Thermo Fisher Scientific) for 1 hour. Spheroids were rinsed twice with warm 1X PBS, treated with 1 mL of complete cortical media per well, and transferred from the agarose microwells into 35 mm-diameter Fluorodishes (World Precision Instruments) containing 1 mL of warm complete cortical media. To examine the effects of centrifugation-based injury in neuronal microtubule structures, we injured another subset of spheroids as described above and fixed overnight with 4% v/v paraformaldehyde and 8% w/v sucrose solution either at 0, 2, 8, and 24 hours post-injury. Following fixation, spheroids were transferred from the agarose wells into 1.5-mL Eppendorf centrifuge tubes, permeabilized and blocked using a 10% bovine serum albumin, 10% normal goat serum and 1% Triton 100-X for 2 hours. Then they were immunolabeled for β_3_-tubulin by treating them with mouse anti-β_3_-tubulin (BioLegend) primary antibodies at a 1:200 dilution overnight at room temperature. The following day, the spheroids were labeled Alexa Fluor 488-conjugated goat anti-mouse secondary antibodies (Jackson Laboratories) at a 1:500 dilution overnight and finally counterstained with 20 µg/mL DAPI solution. Immunolabeled spheroids were transferred to Fluorodishes containing PBS 1X.

### Confocal microscopy

Cortical spheroid images were acquired using an Olympus FV3000 laser scanning confocal microscope at a 30X magnification, oil immersion, 1.5-µm step size in z, and 50 slices per spheroid. 13-29 spheroids per sample were imaged up to 75 microns depth. Images were acquired in Galvano mode. For living spheroid imaging, Fluorodishes containing complete cortical media were kept warm at 37°C by a plate adapter with a built-in thermocouple.

### Image processing and statistical analysis

Images were processed using Fiji v 2.11.0 software Cell viability was determined by counting all EthD1- and Hoechst-labeled nuclei using the 3D Object Counter plug-in (insert reference) and using the following formula to calculate % cell viability:

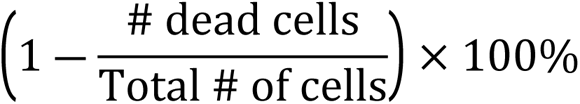

All results are reported as median ± interquartile range in the text. Data visualizations and statistical analysis were performed using GraphPad Prism 9 (GraphPad Software LLC). All data analysis was performed using one-way ANOVA with post-hoc Tukey-HSD tests.

## Results

### Sustained compression injury strains were determined by mechanics modeling and finite element analysis

Injury severity in sustained compression injury is determined by the strains and duration of compression events experienced by the cells in brain tissue, which can be difficult to determine when strain-induced deformation changes cannot be visualized and measured experimentally. In this study, we cannot visualize the strain experienced by the cortical spheroids during centrifugation-induced sustained compression. To estimate said strains, we relied on mechanics modeling and finite element analysis, as detailed in Supplementary Information and (24). We estimated that centrifugation speeds of 209 rads/s and 419 rads/s generated pressures of 38 kPa and 149 kPa, respectively (Fig. 3d). Conversely, we estimated the resulting compressive strains to be -18% and -35%, respectively (Fig. 3b).

### Cortical spheroids showed reduced cell viability following sustained compression

Cellular death is a prominent event following mechanical insults such as sustained compression injury and is responsible for the onset of neuronal loss associated with cognitive loss (26). In many brain injury studies, the injury has been deemed biphasic, with a primary injury of mechanical nature occurring immediately after trauma resulting from the pressures deforming the cells and a secondary injury caused by biochemical signaling triggered by injured cells hours following injury (27). Therefore, we examined the results of sustained compression injury 2, 8, and 24 hours to capture this injury timeline. We found that centrifugation-enabled sustained compression reduced cell viability in cortical spheroids in a strain-dependent manner, with the highest reduction observed 2 hours post-injury (Fig. 4). Spheroids compressed at 38 kPa and 149 kPa were deformed by 18% and 35% strains, respectively. They exhibited significantly lower cell viabilities of 73±10% and 60±24% following 2 hours post-injury when compared to time-matched control spheroids (85±8%, p < 0.0001). Similar viability results were observed at 8 and 24 hours when comparing injured spheroids compressed at 38 kPa and 149 kPa to corresponding time-matched control spheroids. Spheroids compressed at 18% and 35% strains exhibited significantly reduced cell viabilities of 73±8% and 68±10% following 8 hours post-injury and 74±10% and 67±13% following 24 post-injury when compared to those of time-matched control spheroids (84±8% and 91±3%, p < 0.0001). Also, spheroids compressed at 35% strain had lower cell viability than those compressed at 18% strain 2 hours post-injury (p < 0.01) but exhibited similar cell viability reductions after 8- and 24-hours post-injury. These results suggest that most of the observed reduction in viability occurred early following sustained compression injury, where strain magnitude determined the severity of the observed cell viability loss.

**Fig. 4:**
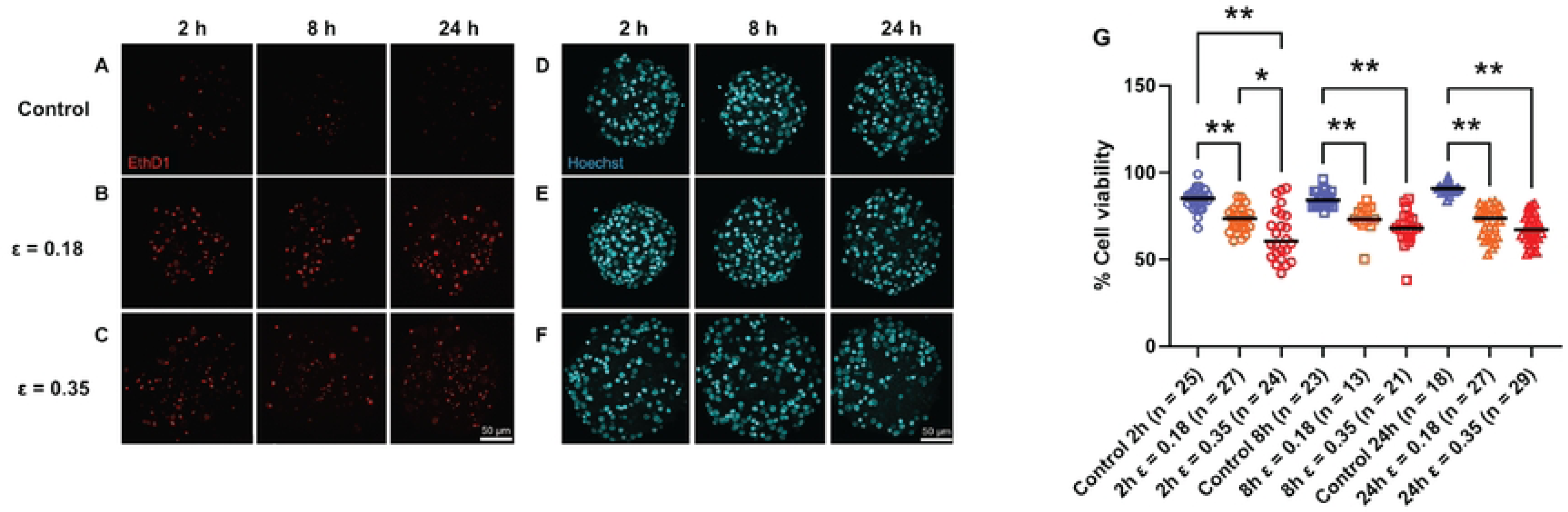
Cell viability loss following sustained compression in spheroids. Cortical spheroids were subjected to sustained compression injury via centrifugation at angular velocities of 209 rad/s (strain of 0.18) and 419 rad/s (strain of 0.35) for 2 minutes. Strains resulting from these angular velocities were determined using mechanics modeling and FEA estimates. (A-F) Living cortical spheroids were labeled for dead cells and total cell nuclei using ethidium homodimer-1 (EthD1, shown in red) and Hoechst 33248 (shown in teal), and imaged 2, 8, and 24 hours post-injury using confocal laser microscopy. (G) Quantification of % cell viability in control and spheroids injured 2, 8, and 24 hours following sustained compression injury. Data reported as median ± interquartile range. Asterisks denote statistical significance (* = *p* < 0.01, ** = *p* < 0.001).

### Cortical spheroids showed disrupted neurite networks following sustained compression

Secondary injury resulting from compressive brain trauma involves a delayed response characterized by biochemical signals that result in neuronal cell loss. Two hours following SCI, cortical microtissues subjected to sustained compression did not exhibit marked differences in neurite network organization and structure when compared qualitatively to control spheroids (Fig. 5). However, 8 and 24 hours after injury, neurite cytoskeletal structures became less defined with more punctate beta-3 tubulin staining, which is associated with microtubule degradation.

**Fig. 5:**
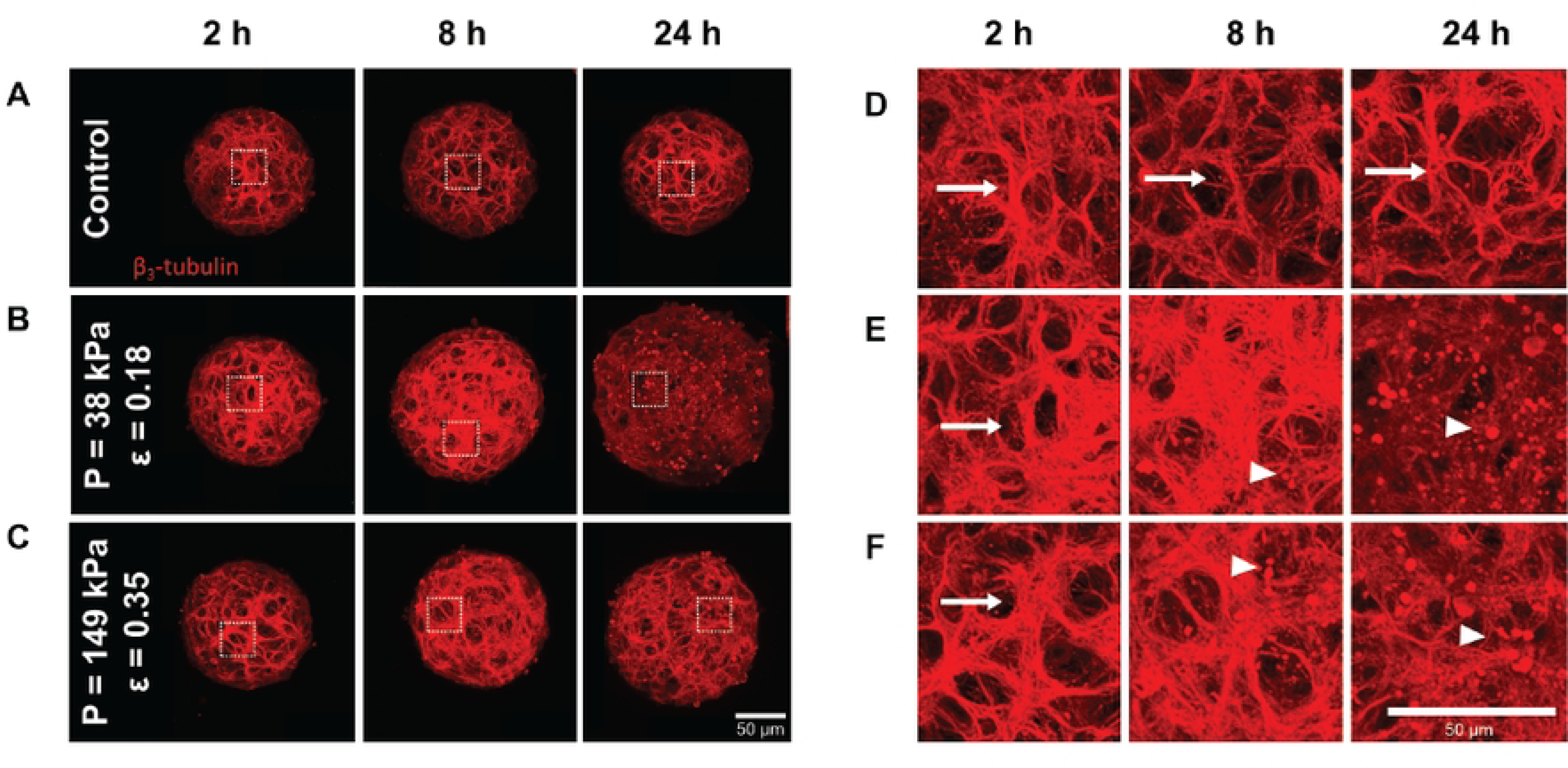
Neurite network disruption in cortical spheroids following sustained compression injury. Confocal z-projections of whole spheroids immunostained for β_3_-tubulin. Cortical spheroids were subjected to sustained compression injury via centrifugation at angular velocities of 209 rad/s (strain of 0.18) and 419 rad/s (strain of 0.35) for 2 minutes. Strains resulting from these angular velocities were determined using mechanics modeling and FEA estimates. Control (A, D) and injured spheroids (B, C, E, F) were fixed 2, 8, and 24 hours post-centrifugation. D, E, F show high magnification views of the corresponding boxed regions in A, B, C. Arrows represent intact microtubule network while arrowheads represent puncta. Scale bars, 50 µm.

## Discussion

Our aims for this study were to highlight a novel centrifugation-based, sustained compression injury modality subjecting cortical spheroids to sustained pressures at low speeds, and to characterize the strains caused by sustained compression using physics-based modeling and finite element analysis. We examined the effects of these strains in cortical neurospheroid cell viability and neurite network organization to observe if sustained pressures achieved via centrifugation would trigger injurious events at the cellular level. Using mechanics-based modeling and FE calculation, we found that strains resulting from sustained compression injury fell in the range of strains reported in mild and moderate brain injury *in vitro* and *in vivo* models (28) as well as from finite element studies (29–31). These strains significantly reduced cell viability and disrupted neuronal cytoskeletal organization in cortical spheroids within the first 24 hours following injury. These results suggest that this injury modality can be used to study sustained compression-based injury while having knowledge of tissue strains. Being able to couple strain information with temporal information on changes in cellular viability and neurite network morphology could provide future insight into injury thresholds and therapeutic time windows following the onset of primary and secondary injury cellular events.

In *in vivo* rodent models, cortical compression has led to increased oxidative stress, astrocyte reactivity, and blood-brain barrier permeability (15), and neuronal hyperexcitability (14). Other *in vitro* studies of sustained compression have observed increased lactate dehydrogenase in response to pressurized 2D cell cultures with neuronal and astrocyte cell lines (16,17). In the present study, we showed strain-dependent decreased cell viability and disrupted neurite network organization in 3D multicellular cortical spheroids. This sustained compression injury experimental modality compressed all spheroids in the same plane while suspended in liquid media. No natural or synthetic hydrogel was used to surround the spheroids, as can be required in some brain injury experiments. The spheroids contained all the key cell types of the brain, including glia; the cells and neurites were interconnected; they produced their own extracellular matrix; and the neurons were electrically active, with mature synapses (20,32). We have recently demonstrated the utility of these cortical spheroids for modeling disease and injury effects, as they can respond to oxygen deprivation and ischemia (33). While the spheroid does not provide the layer-specific organization of the *in vivo* cortex, it is an efficient and importantly, a high-throughput approach that presents many key *in vivo* characteristics. Ongoing studies are examining the responses of the spheroid’s other cell types to sustained compression injury, to leverage the model’s complexity.

Using the centrifuge to study sustained compression injury has several advantages. The spheroids are all likely to experience the same order of magnitude of pressures because they are seeded and self-assembled in the same plane. There are no concerns for anisotropy and heterogeneity in the liquid media. The fluid’s homogeneity and incompressibility facilitate accurate quantification of the pressures experienced by the spheroids. Finally, the centrifuge modality of injury allows for precise control of the pressures by varying the angular velocity and the height of the liquid column.

Our approach also has current limitations. The most prominent is that our calculations for the spheroid strains are likely to be only a first order estimate, for the following reasons. We model the spheroid as a (a) homogenous solid composed of an (b) incompressible (c) neo-Hookean elastic material. Regarding assumption (a): A solid can be assumed to be homogenous if the length-scale at which it first starts being significantly heterogeneous is small compared to the other length scales in the problem. The length scale at which the spheroid starts being significantly heterogeneous (see Fig. 5) is not that much smaller than its overall size. Regarding (b): All materials are compressible at appropriate pressures. Finally, regarding (c): The spheroid’s mechanical behavior is likely to have viscous, plastic, and poro-elastic components in addition to an elastic component. Hence, there are reasons to expect that our assumptions (a)–(c) do not hold perfectly in the experiments, and consequently that our calculations for the strains only provide first order estimates for them. Despite its limitations, our approach is very valuable, because of our approach’s several advantages described above. Further, we note that the estimates for the strains in our approach are only expected to improve in future SCI experiments, as the mechanical characterization of the spheroids becomes optimized.

## Conclusion

In this work we have shown that for sustained compressive injury, the slow loading times involved during centrifugation resulted in low strain rates but still caused significant cellular damage. In contrast to the present work, other studies of brain injury often apply pressures for very short durations at high speeds, and strains and especially strain rates predominantly affect the injury severity (9,10). Our results suggest that in SCI, the deformation, or strain magnitude, is a predominant factor correlating with SCI severity.

The centrifuge modality of injury that we present in this paper is novel, and it can be expanded into a versatile platform for *in vitro* micro-tissue injury studies. It has high throughput, based on the hundreds of spheroids that can be simultaneously loaded. It lends itself to being easily modified so that other more sophisticated pressure time histories can be applied. Additionally, it can be used to study the effect of higher strain rates using more rapidly changing pressures. Finally, we believe that the ease of operation and accessibility of a lab-grade centrifuge, as well as the direct and simple way in which the pressures can be controlled during centrifugation, will make this injury modality relevant to future *in vitro* neurotrauma studies.

## Acknowledgments

We thank Geoffrey Williams for assistance with confocal microscopy.

## References

1. Marmarou A, Fatouros PP, Barzó P, Portella G, Yoshihara M, Tsuji O, et al. Contribution of edema and cerebral blood volume to traumatic brain swelling in head-injured patients. Journal of Neurosurgery. 2000 Aug 1;93(2):183–93.

2. Seano G, Nia HT, Emblem KE, Datta M, Ren J, Krishnan S, et al. Solid stress in brain tumours causes neuronal loss and neurological dysfunction and can be reversed by lithium. Nat Biomed Eng. 2019 Mar;3(3):230–45.

3. Esquenazi Y, Lo VP, Lee K. Critical Care Management of Cerebral Edema in Brain Tumors. J Intensive Care Med. 2017 Jan 1;32(1):15–24.

4. Kochanek PM, Tasker RC, Carney N, Totten AM, Adelson PD, Selden NR, et al. Guidelines for the Management of Pediatric Severe Traumatic Brain Injury, Third Edition: Update of the Brain Trauma Foundation Guidelines. Pediatric Critical Care Medicine. 2019 Mar;20(3S):S1.

5. Hall ED, Bryant YD, Cho W, Sullivan PG. Evolution of Post-Traumatic Neurodegeneration after Controlled Cortical Impact Traumatic Brain Injury in Mice and Rats as Assessed by the De Olmos Silver and Fluorojade Staining Methods. Journal of Neurotrauma. 2008 Mar;25(3):235–47.

6. Clark RS b., Schiding JK, Kaczorowski SL, Marion DW, Kochanek PM. Neutrophil Accumulation After Traumatic Brain Injury in Rats: Comparison of Weight Drop and Controlled Cortical Impact Models. Journal of Neurotrauma. 1994 Oct;11(5):499–506.

7. Kumar A, Alvarez-Croda DM, Stoica BA, Faden AI, Loane DJ. Microglial/Macrophage Polarization Dynamics following Traumatic Brain Injury. J Neurotrauma. 2016 Oct 1;33(19):1732–50.

8. Dalgard C, Cole J, Kean W, Lucky J, Sukumar G, McMullen D, et al. The cytokine temporal profile in rat cortex after controlled cortical impact. Frontiers in Molecular Neuroscience [Internet]. 2012 [cited 2023 Jul 23];5. Available from: https://www.frontiersin.org/articles/10.3389/fnmol.2012.00006

9. Bar-Kochba E, Scimone MT, Estrada JB, Franck C. Strain and rate-dependent neuronal injury in a 3D in vitro compression model of traumatic brain injury. Sci Rep. 2016 Aug 2;6(1):30550.

10. Cullen DK, Vernekar VN, LaPlaca MC. Trauma-Induced Plasmalemma Disruptions in Three-Dimensional Neural Cultures Are Dependent on Strain Modality and Rate. J Neurotrauma. 2011 Nov;28(11):2219–33.

11. Liaudanskaya V, Chung JY, Mizzoni C, Rouleau N, Berk AN, Wu L, et al. Modeling Controlled Cortical Impact Injury in 3D Brain-Like Tissue Cultures. Adv Healthc Mater. 2020 Jun;9(12):e2000122.

12. Shoemaker AR, Jones IE, Jeffris KD, Gabrielli G, Togliatti AG, Pichika R, et al. Biofidelic dynamic compression of human cortical spheroids reproduces neurotrauma phenotypes. Disease Models & Mechanisms. 2021 Dec 22;14(12):dmm048916.

13. Tang-Schomer MD, White JD, Tien LW, Schmitt LI, Valentin TM, Graziano DJ, et al. Bioengineered functional brain-like cortical tissue. Proc Natl Acad Sci U S A. 2014 Sep 23;111(38):13811–6.

14. Ding MC, Wang Q, Lo EH, Stanley GB. Cortical Excitation and Inhibition following Focal Traumatic Brain Injury. J Neurosci. 2011 Oct 5;31(40):14085–94.

15. Lin JL, Huang YH, Shen YC, Huang HC, Liu PH. Ascorbic acid prevents blood-brain barrier disruption and sensory deficit caused by sustained compression of primary somatosensory cortex. J Cereb Blood Flow Metab. 2010 Jun;30(6):1121–36.

16. Murphy EJ, Horrocks LA. A model for compression trauma: pressure-induced injury in cell cultures. J Neurotrauma. 1993;10(4):431–44.

17. Zhang S, Yu R, Zhang Y, Chen K. Cytoprotective effects of urinary trypsin inhibitor on astrocytes injured by sustained compression. Mol Biol Rep. 2014 Mar 1;41(3):1311–6.

18. Boutin ME, Kramer LL, Livi LL, Brown T, Moore C, Hoffman-Kim D. A three-dimensional neural spheroid model for capillary-like network formation. J Neurosci Methods. 2018 Apr 1;299:55–63.

19. Dingle YTL, Boutin ME, Chirila AM, Livi LL, Labriola NR, Jakubek LM, et al. Three-Dimensional Neural Spheroid Culture: An In Vitro Model for Cortical Studies. Tissue Eng Part C Methods. 2015 Dec;21(12):1274–83.

20. Sevetson JL, Theyel B, Hoffman-Kim D. Cortical spheroids display oscillatory network dynamics. Lab Chip. 2021 Nov 25;21(23):4586–95.

21. Rahaman MM, Fang W, Fawzi AL, Wan Y, Kesari H. An accelerometer-only algorithm for determining the acceleration field of a rigid body, with application in studying the mechanics of mild traumatic brain injury. Journal of the Mechanics and Physics of Solids. 2020 Oct 1;143:104014.

22. Wan Y, Fawzi AL, Kesari H. Determining rigid body motion from accelerometer data through the square-root of a negative semi-definite tensor, with applications in mild traumatic brain injury. Computer Methods in Applied Mechanics and Engineering. 2022 Feb 15;390:114271.

23. Wan Y, Fang W, Carlsen RW, Kesari H. A finite rotation, small strain 2D elastic head model, with applications in mild traumatic brain injury. Journal of the Mechanics and Physics of Solids. 2023 Oct 1;179:105362.

24. Wan Y, González-Cruz RD, Hoffman-Kim D, Kesari H. A mechanics theory for the exploration of a high-throughput, sterile 3D in vitro traumatic brain injury model [Internet]. arXiv; 2023 [cited 2023 Aug 31]. Available from: http://arxiv.org/abs/2308.13409

25. Boulet T, Kelso ML, Othman SF. Microscopic magnetic resonance elastography of traumatic brain injury model. Journal of Neuroscience Methods. 2011 Oct 15;201(2):296–306.

26. Xiong Y, Mahmood A, Chopp M. Animal models of traumatic brain injury. Nat Rev Neurosci. 2013 Feb;14(2):128–42.

27. Jassam YN, Izzy S, Whalen M, McGavern DB, El Khoury J. Neuroimmunology of Traumatic Brain Injury: Time for a Paradigm Shift. Neuron. 2017 Sep 13;95(6):1246–65.

28. MacManus DB, Ghajari M. Material properties of human brain tissue suitable for modelling traumatic brain injury. Brain Multiphysics. 2022 Jan 1;3:100059.

29. Viano DC, Casson IR, Pellman EJ, Zhang L, King AI, Yang KH. Concussion in Professional Football: Brain Responses by Finite Element Analysis: Part 9. Neurosurgery. 2005 Nov;57(5):891.

30. Zhang L, Yang KH, King AI. A proposed injury threshold for mild traumatic brain injury. J Biomech Eng. 2004 Apr;126(2):226–36.

31. Carlsen RW, Fawzi AL, Wan Y, Kesari H, Franck C. A quantitative relationship between rotational head kinematics and brain tissue strain from a 2-D parametric finite element analysis. Brain Multiphysics. 2021 Jan 1;2:100024.

32. Dingle YTL, Boutin ME, Chirila AM, Livi LL, Labriola NR, Jakubek LM, et al. Three-Dimensional Neural Spheroid Culture: An In Vitro Model for Cortical Studies. Tissue Eng Part C Methods. 2015 Dec 1;21(12):1274–83.

33. McLaughlin RM, Top I, Laguna A, Hernandez C, Katz H, Livi LL, et al. Cortical Spheroid Model for Studying the Effects of Ischemic Brain Injury. In vitro models. 2023 Apr 1;2(1):25– 41.

